# Prokaryotic mechanosensitive channels mediate copper influx

**DOI:** 10.1101/2025.03.24.644891

**Authors:** Yara Ghnamah, Caitlin D. Palmer, Nurit Livnat-Levanon, Moti Grupper, Amy C. Rosenzweig, Oded Lewinson

**Author notes:** Equal contribution (Yara Ghnamah, Caitlin D. Palmer).

## Abstract

Copper is an essential micronutrient in all kingdoms of life, requiring a meticulous balance between acquisition and toxic overload. While copper import in eukaryotes has been investigated extensively, few prokaryotic copper importers have been identified, leading to the notion that cytoplasmic copper uptake is unnecessary in prokaryotes. Here we report that mechanosensitive channels are key players in prokaryotic copper import. Deletion of the gene encoding the *E. coli* small mechanosensitive channel, _Ec_MscS, leads to significantly reduced copper influx. Conversely, overexpression of _Ec_MscS leads to increased copper influx, elevated intracellular copper content, and renders cells hypersensitive to copper. Furthermore, specific channel blockers and competing permeating ions inhibit _Ec_MscS copper conductance, lowering intracellular copper accumulation and alleviating copper hypersensitivity. These findings extend beyond *E. coli*, as other prokaryotic small mechanosensitive channels of bacterial and archaeal origin also facilitate copper influx. Taken together, these results uncover a previously unknown moonlighting function for mechanosensitive channels as a pathway for prokaryotic copper uptake.

## Introduction

Copper is a vital micronutrient for all organisms, serving as an essential cofactor in critical processes such as oxidative phosphorylation, oxidative stress defense, neurotransmitter synthesis, and iron metabolism^1^. However, excessive copper poses a significant threat, inducing oxidative damage and disrupting iron-sulfur clusters^2,3^. Consequently, organisms have evolved intricate copper handling mechanisms to maintain a precise balance between essential acquisition and potential toxic overload^3–5^. In eukaryotes, copper is first imported into the cytoplasm by transporters of the CTR family^6^. Subsequently, it is directed to specific cellular compartments through the coordinated activity of copper chaperones and additional transporters. In cases of copper overload, ATP-driven copper efflux pumps are redirected from membranes of intracellular compartments to the cell surface, thereby averting its toxic effects^7^. There are many similarities between prokaryotic copper efflux proteins and their eukaryotic counterparts. For example, CopA, the *E. coli* P-type ATPase copper efflux pump, exhibits 33% sequence identity and 61% sequence similarity with its human equivalents ATP7A and ATP7B which mediate copper efflux and cuproproteins metalation, respectively ^8,9^, and the structure of the archetypical yeast copper chaperone Atx1 closely mirrors that of its bacterial homologues ^10,11^.

In contrast, with a few rare exceptions^12–14^, bacterial counterparts to the eukaryotic high affinity copper importers have not yet been identified, leading to the accepted notion that bacterial copper importers may not exist. Nevertheless, copper must enter the prokaryotic cytoplasm, as evidenced by the prevalence of copper-responsive regulatory elements in bacterial genomes, some of which respond to picomolar copper concentration^15^. Moreover, the metalation of certain cuproproteins has been demonstrated to follow a mechanism requiring the delivery of copper first to the cytoplasm^12^, followed by metalation through copper efflux to the periplasm. Given the charge density and extremely low membrane permeability of copper, its entry by unmediated passive diffusion is highly improbable. As such, the mechanism facilitating copper transport to the prokaryotic cytoplasm remains one of the most significant conundrums in biological metal homeostasis.

One way to identify gene functions or find moonlighting functions of known genes is phenomic profiling. In a comprehensive study by Nichols et al., the phenotypes of thousands of *E. coli* single gene deletion mutants under hundreds of growth conditions were assessed ^16^. One curious observation from these data is that a strain lacking the small mechanosensitive channel (_Ec_MscS) exhibited an increased tolerance to high copper concentrations. The primary function of mechanosensitive channels is to maintain cell turgor in prokaryote^17–19^. Under hypoosmotic conditions, water enters prokaryotic cells, causing cell swelling. Mechanosensitive channels respond to the increased membrane lateral pressure and open rapidly, allowing the passive diffusion of ions and small solutes out of the cell and preventing cell rupture^20^. Given its established function as a channel, we hypothesized that the copper tolerance phenotype of the _Ec_MscS deletion strain could be due to a secondary role in copper permeation.

Herein, we demonstrate that mechanosensitive channels indeed play a role in mediating copper influx. We show that deletion of the gene encoding _Ec_MscS leads to reduced intracellular copper content, while its overexpression results in increased copper influx and copper sensitivity. Furthermore, observed copper sensitivity can be alleviated by the addition of competing ions or channel specific blockers. Finally, similar results were obtain using six _Ec_MscS homologues from bacteria and archaea, suggesting that this mechanism of copper import is widespread in prokaryotes. Mechanosensitive channels are thus versatile regulators of both osmotic and copper homeostasis.

## Results

To investigate the potential involvement of _Ec_MscS in copper influx, we first employed a gene reporter approach. Specifically, we utilized the promoter region of the gene encoding the copper efflux pump CopA, whose activity is tightly regulated by intracellular copper levels^21^. This promoter can therefore serve as an indicator of internal copper concentration^22^. We cloned the *copA* promoter region from the genome of wildtype (WT) *E. coli* MG1655 and fused it upstream of the *lux* operon genes (*luxCDABE*), creating the luminescence reporter plasmid *pcopA-Lux*. Next, we transformed both the WT strain and its isogenic _Ec_MscS deletion strain (hereafter referred to as Δ*mscS*) with the *pcopA-Lux* reporter plasmid.

Cells were grown in minimal media to mid-exponential phase, adjusted to identical densities, and transferred to a microplate reader. Following equilibration, 100 μM Cu²⁺ (as CuSO₄) was injected into the cell suspension. As shown in Figure 1A, consistent with our hypothesis that _Ec_MscS mediates copper permeation, the rate of entry and steady state accumulation of copper was significantly lower in Δ*mscS* cells than in WT cells. To further establish the link between _Ec_MscS expression and copper permeation, we lowered the osmolarity of the growth medium (see Methods for full details), a condition known to increase the expression of endogenous _Ec_MscS^23^. Before proceeding with activity measurements, we verified that the viability of WT and Δ*mscS* cells was not compromised under these conditions (Figure S1). As shown in Figure 1B, consistent with the proposed role of _Ec_MscS in mediating copper influx, exposure of WT cells to mild hypoosmotic conditions resulted in increased luminescence, indicative of elevated copper entry. This effect appeared specific to cells expressing _Ec_MscS, as the luminescence recorded in Δ*mscS* cells remained unaffected by exposure to mild hypoosmotic conditions.

**Figure 1.**
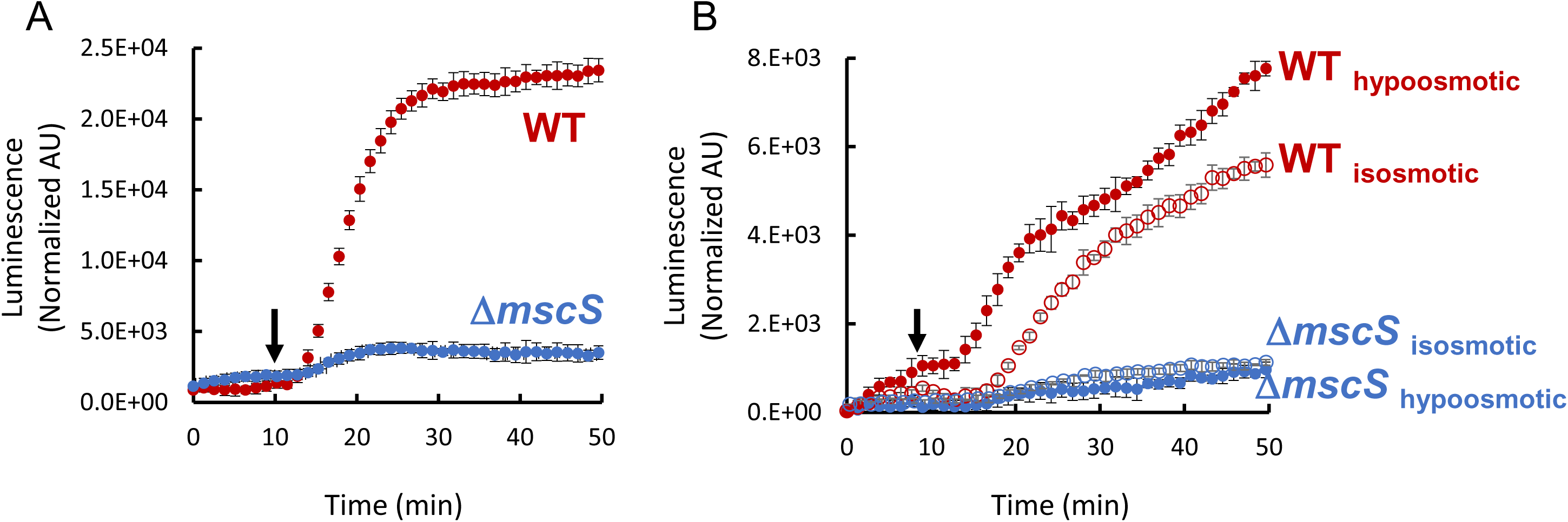
_EC_MscS facilitates copper permeation. (A) At the time indicated by the arrow, 100 mM CuSO_4_ was injected to cultures of WT or *ΔmscS* cells (red and blue curves, respectively) expressing the copper reporter *pcopA-Lux.* Luminescence was normalized by the optical density of the cultures. (B) Same as A, only the cells were pre-exposed to isosmotic or mild hypoosmotic conditions (open and closed circles, respectively. See methods for full details). Results are the mean of biological triplicates, with error bars indicating standard deviations.

Next, we further assessed intracellular copper levels via copper sensitivity growth assays. As mentioned above, cells lacking the gene encoding _Ec_MscS are more tolerant of elevated (and thus toxic) concentrations of environmental copper ^16^. Accordingly, we reasoned that WT cells overexpressing _Ec_MscS would display increased sensitivity to copper. To overexpress _Ec_MscS in WT *E. coli* cells, we used the same pET-28 MscS construct used for its original structural deteremination^24^. In the absence of copper, cell growth was not affected by overexpression of _Ec_MscS (Figure 2A), demonstrating that under these conditions, _Ec_MscS expression itself does not pose a significant fitness cost. We then grew the cells at high copper concentrations and observed that, across a broad concentration range and in both complex and defined media, cells expressing _Ec_MscS exhibited pronounced hypersensitivity to copper. This hypersensitivity strictly depended on the presence of the expression inducer (IPTG), further linking _Ec_MscS to copper permeation (Figure 2A-B, Supplementary Figure S2). Because *E. coli* is inherently quite resilient to environmental copper^3^, these copper sensitivity assays described above required usage of millimolar copper concentrations, raising questions about their physiological relevance.

**Figure 2.**
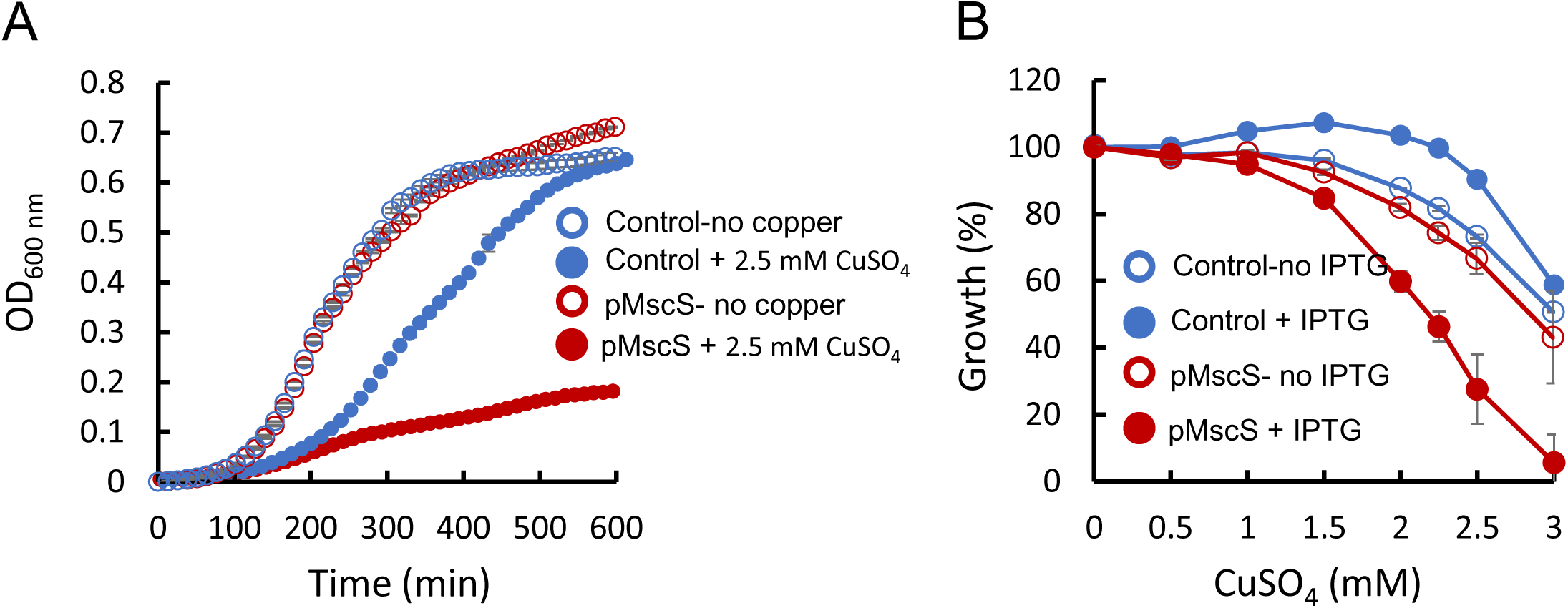
_EC_MscS overexpression increases copper sensitivity. (A) Cells transformed with an empty control vector or the _EC_MscS expression vector pMscS (blue and red curves, respectively) were cultured in the absence or presence of 2.5 mM CuSO_4_ (open and closed symbols, respectively). Optical density at 600 nm was measured every 5 min. (B) Cells transformed with an empty control vector or the _EC_MscS expression vector pMscS (blue and red curves, respectively) were cultured for 10 h in the presence of the indicated concentration of CuSO_4_ in the absence or presence of 0.01 mM IPTG (open and closed symbols, respectively). Growth in the absence of cooper was defined as 100%. Averages of biological triplicates are shown, with error bars indicating standard deviations (shown unless smaller than icons).

To investigate copper permeation more directly and at lower copper concentrations, we employed inductively coupled plasma mass spectrometry (ICP-MS) to monitor intracellular copper content of cells overexpressing _Ec_MscS. Cultures were grown to mid-exponential phase in the presence of the inducer IPTG, followed by the addition of micromolar concentrations of copper. After a 30 min incubation, a sample was withdrawn for analysis. To terminate the reaction, cells were pelleted, washed three times with an EDTA-containing buffer, and their intracellular copper content was measured using ICP-MS. As shown in Figure 3, the overexpression of _Ec_MscS resulted in up to a 10-fold increase in intracellular copper content. These findings strongly suggest that _Ec_MscS-mediated copper permeation occurs at physiological copper concentrations. However, an alternative explanation could be that the expression and/or activity of _Ec_MscS compromises membrane barrier function, leading to nonspecific leakage.

**Figure 3.**
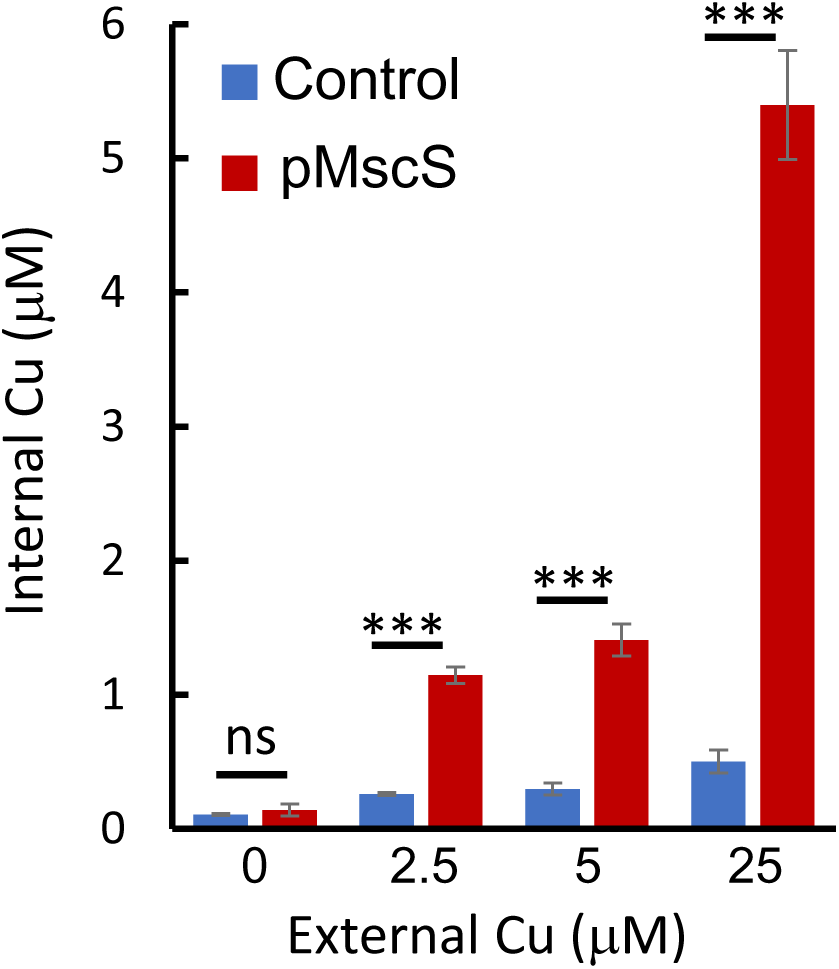
_EC_MscS expression increases intracellular copper accumulation. (A) Cells transformed with an empty control vector or the _EC_MscS expression vector pMscS (blue and red bars, respectively) were cultured in the presence of IPTG (0.01 mM) to mid log phase and then exposed for 30 min to the indicated concentrations of CuSO_4_. Cells were washed with an EDTA containing buffer and their intracellular copper concentration was deteremiend by ICP-MS. Shown are averages of biological triplicates, with error bars indicating standard deviations. *** p < 0.005; ns, not significant, as determined by a two-tailed Student’s t-test.

To address this possibility, we evaluated the impact of _Ec_MscS on cell sensitivity to various agents, including compounds that act directly on the membrane (e.g., detergents, ethanol, EDTA) and those requiring intracellular permeation to exert antibacterial effects (e.g., norfloxacin, tetracycline). As illustrated in Supplementary Figure 3, the overexpression of _Ec_MscS had no impact on sensitivity to any of the tested agents. This suggests that _Ec_MscS overexpression does not alter membrane permeability, supporting the conclusion that the observed copper sensitivity and accumulation specifically result from copper permeation via _Ec_MscS.

Next, we leveraged the robustness and high throughput of the copper sensitivity assays (Figure 2) to explore additional conditions under which _Ec_MscS activity is expected to either increase or decrease. To simulate increased _Ec_MscS activity, we utilized the A106V mutant, which is known to stabilize an open subconducting state of _Ec_MscS and preferentially conduct divalent ions such as Ca²⁺ and Ba²⁺ ^25,26^. In the absence of copper, the expression of the A106V variant had no discernible effect on cell growth (Figure 4A). However, in the presence of copper, cells expressing the A106V variant exhibited greater copper sensitivity compared to those expressing WT _Ec_MscS (Figure 4B), consistent with the proposed role of _Ec_MscS in copper conductance,.

**Figure 4.**
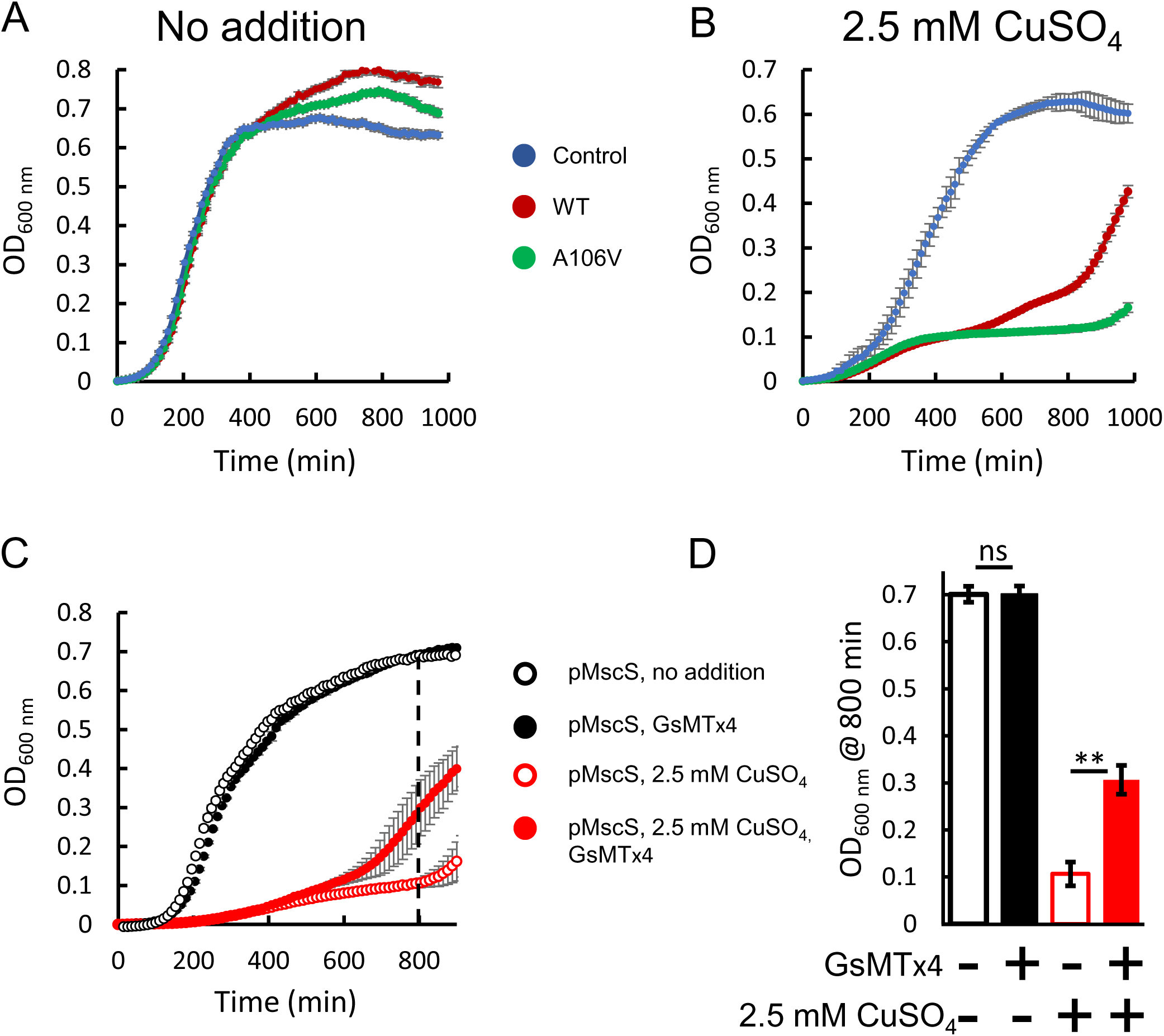
Copper sensitivity correlates with the activity level of _EC_MscS. Cells transformed with an empty control vector or vectors encoding _Ec_MscS WT or A106V (blue, red, and green curves, respectively, as indicated) were cultured in the absence (A) or presence (B) of 2.5 mM CuSO_4_. (C) Cells transformed with a vector encoding WT _EC_MscS (all curves) were cultured in the absence or presence of 2.5 mM CuSO_4_ (black and red curves, respectively) and in the absence or presence of 2 mM GsMTx4 inhibitor (open and closed symbols, respectively). The dashed line represents data points used to generate the bar graph shown in D. (D) Optical density (OD_600_) at 800 min (dashed vertical line in C) of cells grown in the absence or presence of 2.5 mM CuSO_4_ and in the absence (−) or presence (+) of 2 mM GsMTx4 inhibitor, as indicated. Results are averages of biological triplicates with error bars indicating standard deviations (shown unless smaller than the icons). ** p < 0.02; ns, not significant, as determined by a two-tailed Student’s t-test.

To mimic conditions of reduced _Ec_MscS activity, we expressed _Ec_MscS in the presence of the tarantula venom-derived peptide GsMTx4, which is a non-competitive partial blocker of _Ec_MscS, reducing its open probability (P_o_)^27^. In the absence of copper, the GsMTx4 inhibitor showed no impact on the growth of _Ec_MscS-expressing cells (Figure 4C). However, in the presence of copper, the GsMTx4 inhibitor partially alleviated the _Ec_MscS-mediated copper sensitivity (Figure 4C-D). Importantly, this partial alleviation of copper sensitivity aligns well with the the reported inhibitory potency of ∼25% of GsMTx4^28^.

As an additional strategy to reduce the apparent _Ec_MscS-mediated copper uptake, we employed competing ions. _Ec_MscS is known to conduct Li^+^ and Ca^2+ 26^, which are largely nontoxic to bacteria. We posited that the presence of such competing nontoxic ions would reduce copper flux and result in the partial alleviation of its toxic effects. In the absence of copper, Li^+^ and Ca^2+^ had no discernible effect on cell growth. In contrast, in the presence of copper, both Li^+^ and Ca^2+^ partially alleviated the copper sensitivity of _Ec_MscS-expressing cells (Figure 5A). The competitive nature of Li^+^ and Ca^2+^ with copper for permeation via EcMscS was further evident in the intracellular copper content of the cells, as measured by ICP-MS. Both ions significantly reduced the copper content of _Ec_MscS-expressing cells (Figure 5B). In summary, the collective findings from the gain-of-function mutant A106V (Figure 4A-B), the tarantula GsMTx4 inhibitor (Figure 4C-D), and competition with calcium and lithium (Figure 5A-B) strongly support the conclusion that copper influx is indeed mediated by _Ec_MscS.

**Figure 5.**
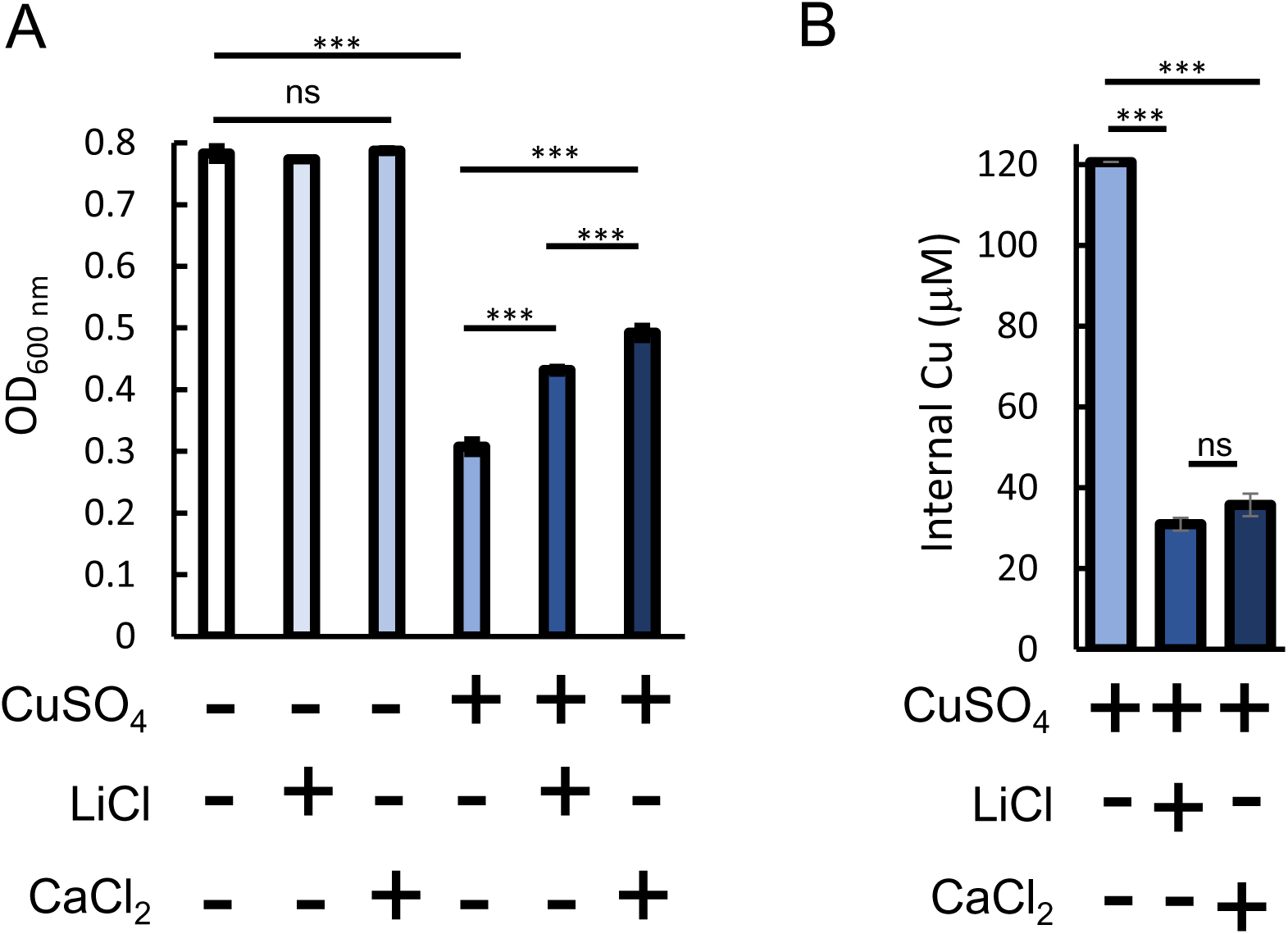
Competing ions reduce _Ec_MscS-mediated copper influx. Cells transformed with the _EC_MscS expression vector pMscS were cultured in LB media in the presence or absence of copper (2.5 mM), lithium (12.5 mM), and calcium (5 mM), as indicated. The optical density at 600 nm was measured after 10 h of growth. (B) Copper content of cells expressing _Ec_MscS exposed for 30 min to the indicated metals (or their combinations). Averages of biological triplicates are shown, with error bars indicating standard errors (shown unless smaller than icons). *** p < 0.001; ns, not significant, as determined by a one-way ANOVA and Tukey HSD.

Compared to selective ion channels (*e.g.,* sodium and potassium ion channels), mechanosensitive channels are relatively nonselective, and any ion with a hydrated radius of 3.2-4.5 Å may potentially permeate^26^. With a hydrated radius of 4.2 Å, Cu^2+^ falls well within this range. However, the hydrated radii of other divalent metal ions, such as Zn^2+^ and Cd^2+^, also fall within this range, suggesting the potential for permeation via _Ec_MscS. To explore this possibility, we conducted metal sensitivity assays with Zn^2+^ and Cd^2+^. As shown in Figure 6A-B, cells overexpressing _Ec_MscS exhibited increased sensitivity to both Cd^2+^ and Zn^2+^, indicating that these metals also permeate via _Ec_MscS. To support these findings, we compared the intracellular content of control and _Ec_MscS-expressing cells after 30 min exposure to either Cd^2+^ or Zn^2+^. Expression of _Ec_MscS led to approximately a 3- and 6-fold increase in intracellular accumulation of zinc and cadmium, respectively (Figure 6C). Thus, while there may be some selectivity for copper, _Ec_MscS can mediate uptake of other metal ions.

**Figure 6.**
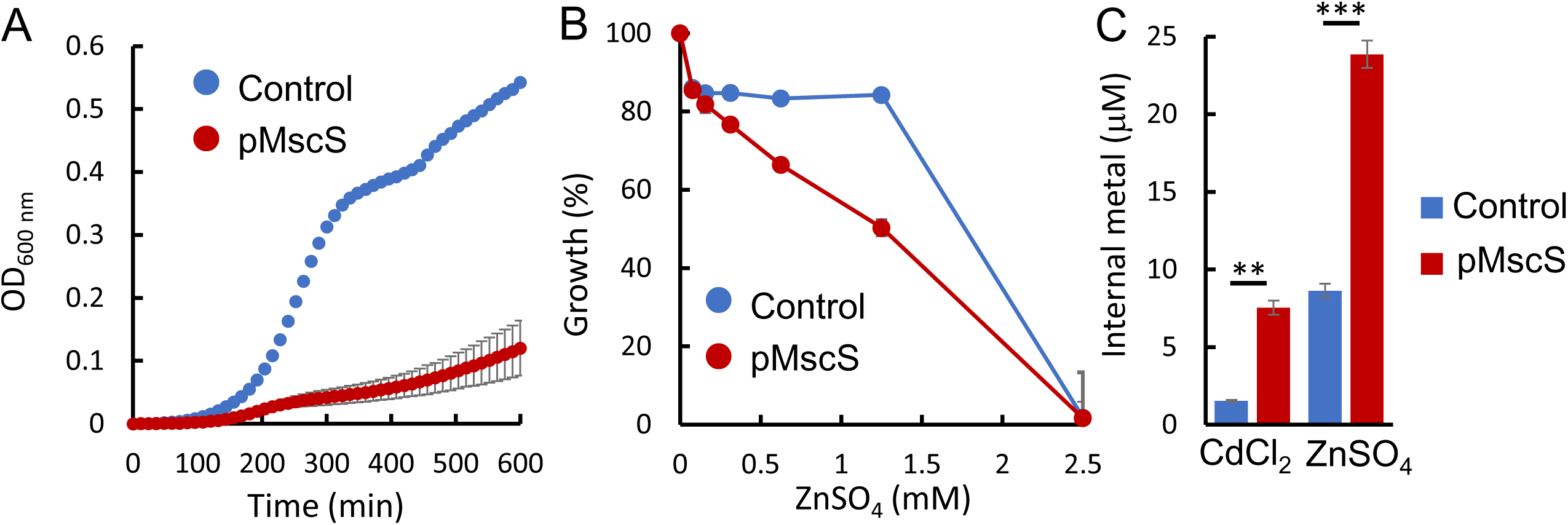
_Ec_MscS mediates influx of additional metals. (A) Cells transformed with an empty control vector or the _EC_MscS expression vector pMscS (blue and red curves, respectively) were cultured in the presence of 250 mM CdCl_2_. Optical density at 600 nm was measured every 5 min. (B) Cells transformed with an empty control vector or the _EC_MscS expression vector pMscS (blue and red curves, respectively) were cultured in the presence of the indicated concentrations of ZnSO_4_, and the optical density at 600 nm was measured after 10 h of growth. Shown is the growth, in percent, relative to the growth in the absence of metal. (C) Metal content of control or _Ec_MscS-expressing cells exposed for 30 min to either 250 mM CdCl_2_ or 1.25 mM ZnSO_4_, as indicated. *** p < 0.001; ** p < 0.005; as determined by a two-tailed Student’s t-test. In A-C, resulst are averages of biological triplicates, with error bars reperesnting standard errors (shown unless smaller than the icons).

Next, we tested whether copper permeation is a unique function of _Ec_MscS or if it is shared by other mechanosensitive channels. Homologs of EcMscS are widely distributed across bacteria and archaea, exhibiting substantial sequence conservation (Figure S4). Given this high level of conservation, we hypothesized that other mechanosensitive channels might also facilitate copper influx, similar to _Ec_MscS. To test this, we transformed WT *E. coli* with expression plasmids encoding four bacterial small mechanosensitive channels (*Listeria monocytogenes*, *Salmonella enterica*, *Thermotoga maritima*, and *Thermus thermophilus*) and two archaeal channels (*Thermoplasma volcanium* and *Archaeoglobus fulgidus*). Cultures were grown to mid-exponential phase, exposed to copper for 30 min, and their intracellular metal content was quantified using ICP-MS. As shown in Figure 7A, expression of the MscS homologs led to a 1.5-fold (*Listeria monocytogenes*) to 8-fold (*Salmonella enterica*) increase in intracellular copper. Accordingly, over a range of concentrations, *E. coli* cells expressing the MscS homologs were more sensitive to copper (Figure 7B). The differences in copper conductance activity between the MscS homologues may stem form different expression levels or from variable compatibility/responsiveness to *E. coli* lipids. Nevertheless, these findings demonstrate that copper conductance is not unique to _Ec_MscS and suggest that this function is likely conserved among prokaryotic small mechanosensitive channels.

**Figure 7.**
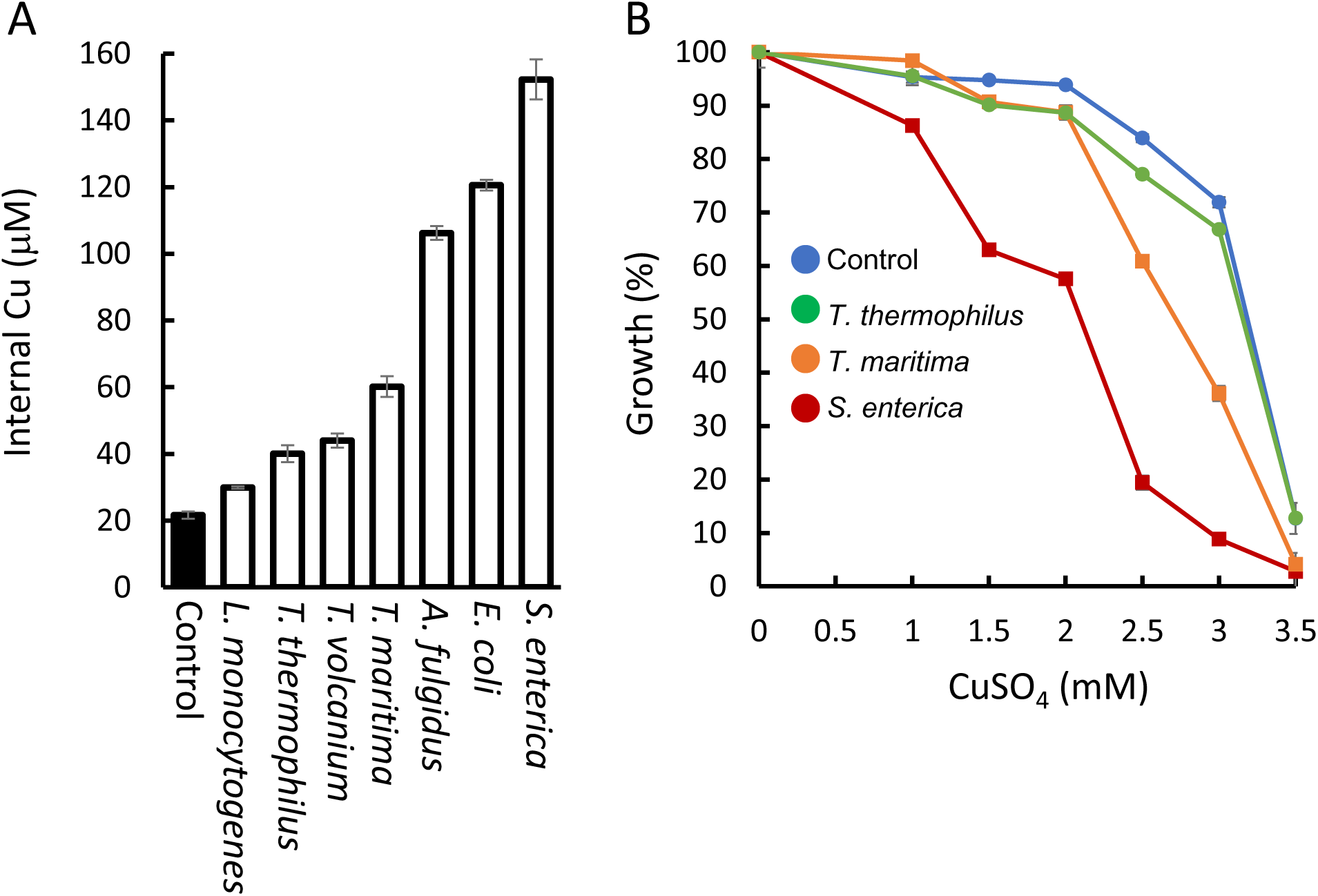
Copper conductance is conserved among prokaryotic mechanosensitive channels. (A) Cells were transformed with an empty control vector or vectors encoding MscS homologues from the indicated organisms, and their intracellular content was determined using ICP-MS following exposure to 2.5 mM CuSO_4_. (B) Cells transformed with an empty control vector (blue curve) or vectors encoding MscS homologues from *Thermus thermophilus, Thermotoga maritima,* or *Salmonella enterica* (green, orange, and red curves, respectively) were cultured in the presence of the indicated concentrations of CuSO_4_, and the optical density at 600 nm was measured after 10 h of growth. Averages of biological triplicates are shown, with some errors bars (indicating standard deviations) not visible as they are smaller than icons.

## Discussion

Eukaryotes and prokaryotes both employ copper-specific efflux pumps to mitigate the toxic effects of copper overload^3,29,30^. However, while copper import systems have been extensively characterized in eukaryotes^6^, equivalent mechanisms in prokaryotes remain poorly understood. This has contributed to the prevailing notion that prokaryotes lack dedicated copper importers. Supporting this perspective is the observation that d-block metals in prokaryotes are imported by transporters belonging to the ATP-binding cassette (ABC) superfamily^31^. Although numerous ABC transport systems for the import of zinc, manganese, iron, cobalt, nickel, and molybdate have been identified across various bacterial species^14,32–37^, a copper-importing ABC transporter does not seem to exist.

One possible explanation for this conundrum is that copper is imported through an alternative mechanism whose primary function is unrelated to metal homeostasis. Here, using a luminescence reporter gene system, direct measurements of intracellular copper by ICP-MS, and copper sensitivity assays, we demonstrate that copper enters *E. coli* cells through the small mechanosensitive channel, _Ec_MscS.

In retrospect, these findings are perhaps not so surprising: small mechanosensitive channels are not very specific, permitting the passage of various monovalent and divalent ions, such as potassium, barium, and calcium^26^. Given copper’s valence of +1 or +2 and its hydrated radius, which is similar to that of barium and calcium, it is reasonable to infer that copper can also permeate through small mechanosensitive channels. We also find that this “moonlighting” activity is not unique to _Ec_MscS, but is also shared by other small prokaryotic mechanosensitive channels of both archaeal and bacterial origin, providing a compelling answer to the longstanding question of how copper is acquired by prokaryotic cells.

## Material and Methods

### Bacterial strains and plasmids

Unless otherwise stated, the *E. coli* wild-type strain BW25113 or its Δ*mscS* isogenic derivative were used in all experiments. To generate the plasmid *_p_copA-Lux*, the genomic region corresponding to the promoter of *copA* (nucleotide positions 511354–511573) was amplified from a single colony of *E. coli* MG1655 using standard PCR. Using restriction free cloning, the amplicon was then inserted into the *_p_PL2lux* plasmid upstream of the *luxCDABE* operon. The pET28-MscS plasmid for expression of _EC_MscS was purchased from the Addgene nonprofit plasmid repository deposited by the Rees group ^38^. The plasmids for expression of the MscS homologues shown in Figure 7 are also pET-28 derivatives and were a kind gift from the Rees group at Caltech.

### Cell growth for luminescence measurements

Cells were transformed with the *_p_copA-Lux* plasmid described above and a single colony was used as inoculum. Cultures were grown aerobically at 37 °C in Citrate-Phosphate Defined Medium (pH 7). This medium has an osmolality of 220 mOsM and contains the following components per liter: 8.58 g Na₂HPO₄, 0.87 g K₂HPO₄, 1.34 g citric acid, 1.0 g (NH₄)₂SO₄, 0.001 g thiamine, 0.1 g MgSO₄·7H₂O, 0.002 g (NH₄)₂SO₄·FeSO₄·6H₂O. After an overnight culture in the presence of 0.04% (w/v) glucose, glucose was supplemented to a final concentration of 0.2% (w/v), and the cultures were grown for another 2-3 hours, after which they were diluted to OD₆₀₀ of 0.1 with identical media supplemented with 0.5 M NaCl. The cultures were then grown to an OD₆₀₀ of 0.3-0.5, at which point the cultures were diluted 20-fold into pre-warmed medium, with or without 0.5 M NaCl for generation of isosmotic or hypoosmotic conditions, respectively.

### Luminescence measurements

For luminescence measurements, the cultures were diluted to an OD₆₀₀ of 0.025 using identical media, and 150 µL were transferred to a covered 96-well plate. The plate was incubated at 37 °C in an automated plate reader (InfinitePro 200, Tecan), with intermittent shaking and continuous monitoring of OD₆₀₀ and luminescence. All experiments were performed in biological triplicates, with at least three technical replicates each.

### Copper sensitivity assays

Cultures were grown in LB media or in M9 minimal media supplemented with 0.5% glycerol and 0.1% tryptone, as indicated. Following ∼16 h incubation with constant agitation (220 rpm, 37 °C) cultures were diluted to an OD_600_ of 0.05 in the absence or presence of 0.1 mM IPTG and the indicated concentrations of CuSO_4_. 150 µL were transferred to a covered 96-well plate. The plate was incubated at 37 °C in an automated plate reader (InfinitePro 200, Tecan), with intermittent shaking and continuous monitoring of OD₆₀₀. All experiments were performed in biological triplicates, with at least three technical replicates each.

### ICP-MS measurements of intracellular metal content

Cultures were grown in the absence of metals at 200 rpm and 37 °C to an OD_600_ of 1.0 in M9 medium supplemented with 50 μg/mL ampicillin, followed by induction with 100 μM IPTG. After incubation for 2.5 h, the OD_600_ values of the cultures were measured, and cell aliquots were placed on ice for 30 min before centrifugation at 4000g for 5 min. Cells were washed with 50 mM potassium phosphate (pH 7.5) and 2 mM MgCl_2_ and resuspended in the same buffer to an OD_600_ of 2.5. After 10 min recovery in the presence of 0.2% glucose at 200 rpm and 37 °C, metals were added at the indicated concentrations. Metals were freshly prepared as 0.100 M stock solutions of CuSO_4_, CdSO_4_, and ZnCl_2_ using MilliQ water and acid treated glassware. After 30 min, a 2-5 mL sample was withdrawn, centrifuged at 4,000g for 5 min, washed with 50 mM potassium phosphate (pH 7.5) and 10 mM EDTA, and centrifuged again for a total of three times. Cell pellets were resuspended in 5 mL of MilliQ water, and metal contents were measured directly by inductively coupled plasma mass spectrometry (ICP-MS, Thermo iCAP Q, Northwestern Quantitiative Biological Imaging Facility). ICP-MS samples were prepared in 5% trace metal free nitric acid and 1% trace metal free hydrogen peroxide with 5 ppb indium, lithium, scandium, and yttrium standards (Inorganic Ventures). The concentrations of iron, cobalt, nickel, copper, cadmium, lithium, calcium, and zinc were measured using the NWU-16 (Inorganic Ventures) multielement standard. Intracellular metal concentrations were calculated using the metal mol amount determined by ICP-MS, the conversion of culture volume and optical density to number of cells using 8×10^8^ cells per 1 mL of OD_600_ of 1^39^, and the estimated *E. coli* cell volume of 1 fL.

### Sequence alignment

Sequence alignment of MscS homologues was performed using the Geneious Prime (2025.0.3) software, using the built-in MUSCLE alignment tool with default parameters.

## Data availability

All data generated or analyzed during this study are included in the published article or Extended Data Information.

## Acknowledgments

This work was supported by NSF-BSF MCB grant 1938715 (A.C.R., O.L.). C. D. P. was supported in part by NIH grant T32GM008382. Research in the Lewinson lab is supported in part by the Rappaport Institute for Biomedical research. ICP-MS analysis was performed at the Quantitative Bio-element Imaging Center at Northwestern University, supported by NASA Ames Research Center Grant NNA04CC36G.

## Author contributions

OL and ACR conceptualized the work, and YG, CDP, NLL, MG designed experiments. YG, CDP, and NLL performed experiments, analyzed data, and prepared figures. YG, CP, NLL, OL, and ACR contributed to writing and editing the manuscript.

