## Supplementary information for "Prokaryotic mechanosensitive channels mediate copper influx"

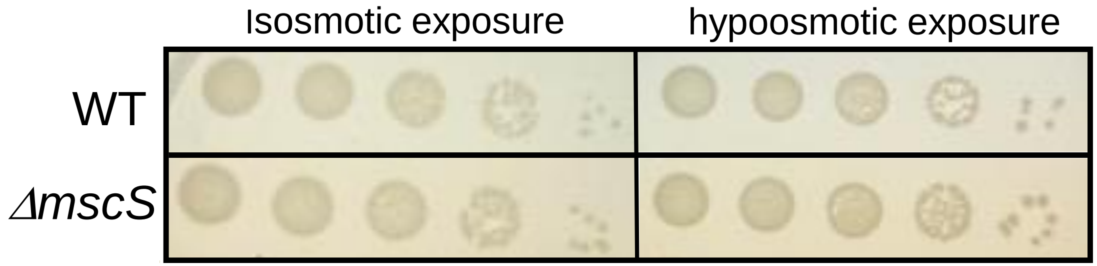


**Supplementary Figure S1. WT and Δ*mscS* cells are equally insensitive to mild hypoosmotic conditions.** (A) Cultures of WT or Δ*mscS* cells (as indicated) were grown to mid exponential phase, diluted to $OD_{600}$ of 0.025, and exposed to isosmotic or hypoosmotic conditions as detailed in the methods section. 4.5 mL were spotted in serial 10-fold dilutions from left to right onto LB-Agar plates.

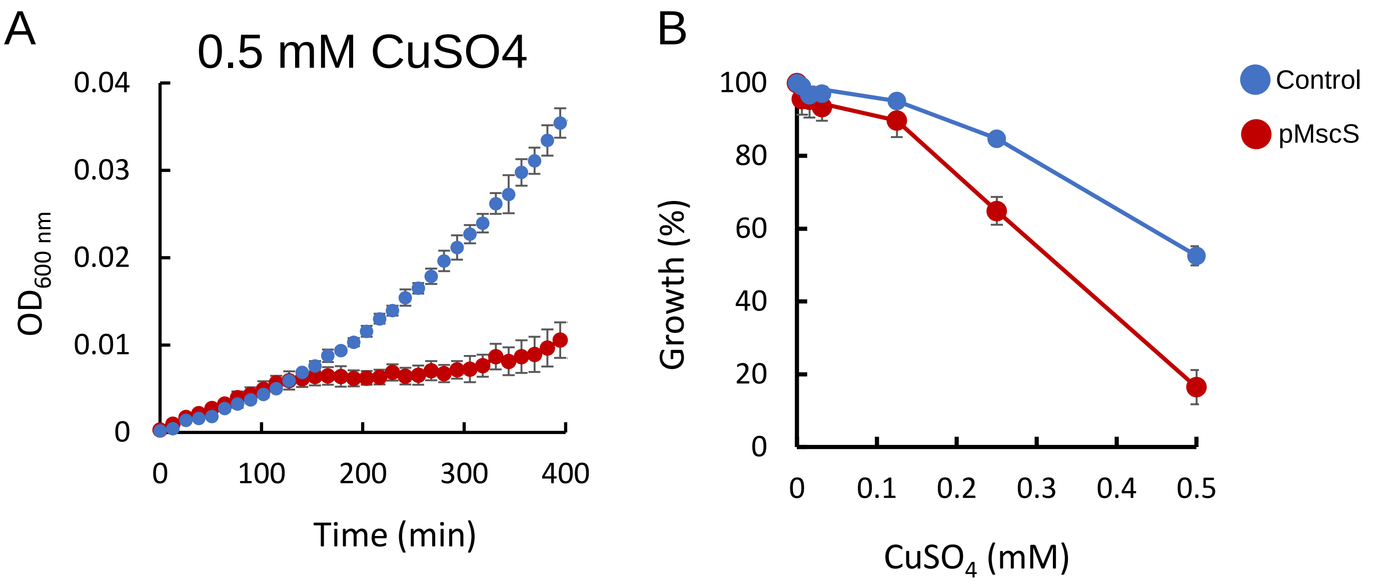


**Supplementary Figure S2. Copper sensitivity in minimal media.** (A) Cells transformed with an empty control vector or the vector encoding $_{EC}$MscS (blue and red curves, respectively) were cultured in M9 minimal media in the presence of 0.5 mM $CuSO_4$. Optical density at 600 nm was measured every 5 min. (B) Cells transformed with an empty control vector or the vector encoding $_{EC}$MscS (blue and red curves, respectively) were cultured in M9 minimal media for 10 h in the presence of the indicated concentration of $CuSO_4$ in the presence of 0.01 mM IPTG. Growth in the absence of cooper was defined as 100%. Averages of biological triplicates are shown, with error bars indicating standard deviations (shown unless smaller than icons).

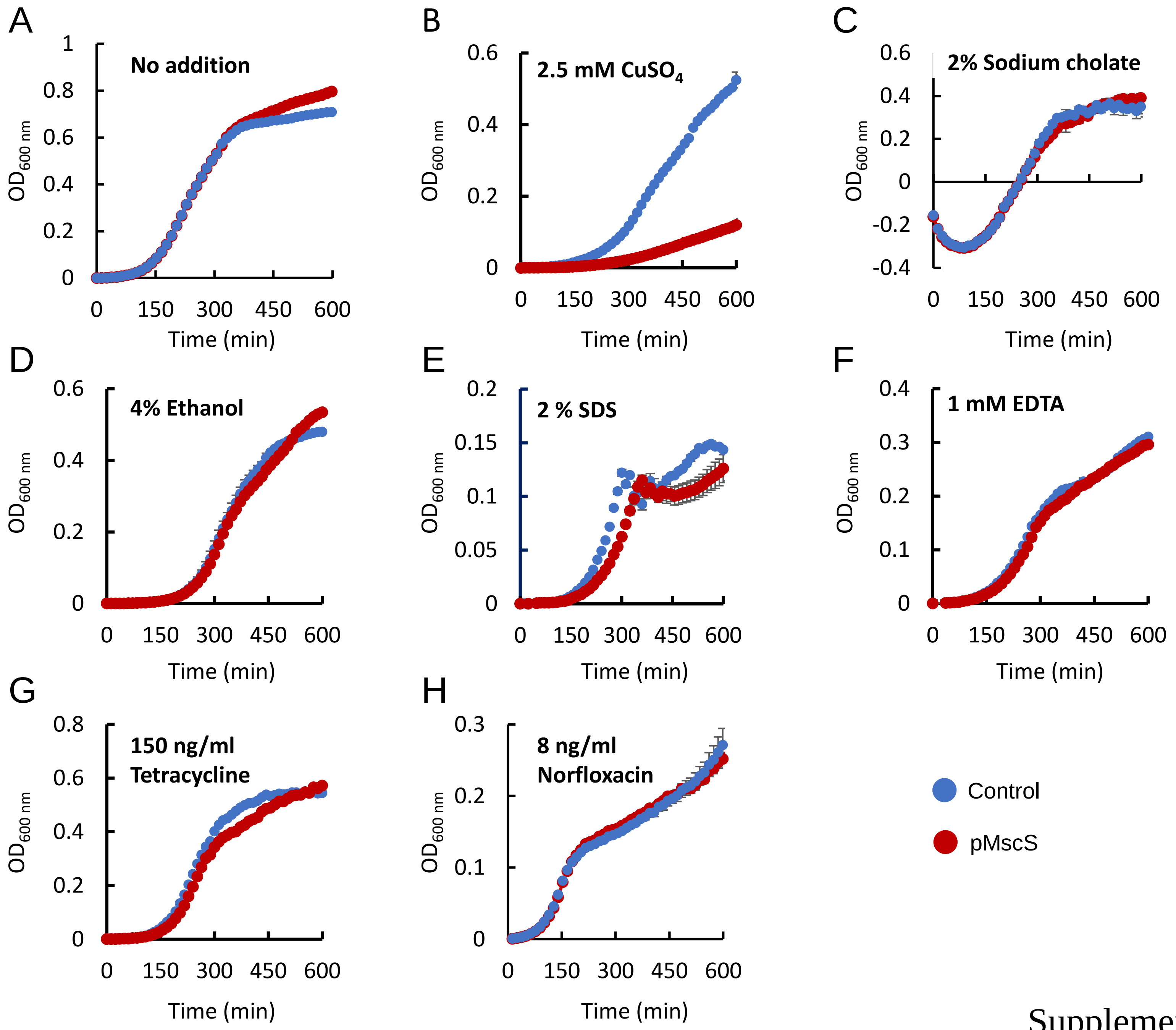


Supplementary Figure 3

**Supplementary Figure S3. Expression of $_{EC}$MscS does not increase cells' sensitivity to a variety of stressors.** Cells transformed with an empty control vector or the vector encoding $_{EC}$MscS (blue and red curves, respectively) were cultured LB media in the presence of 0.01 mM IPTG and in the absence (A) or presence of 2.5 mM $CuSO_4$ (B), 2% sodium cholate (C), 4% ethanol (D), 2 % SDS (E), 1 mM EDTA (F), 150 ng/ml Tetracycline (G), or 8 ng/ml Norfloxacin (H). Optical density at 600 nm was measured every 5 min, and results are averages of biological triplicates. Error bars, indicating standard deviations are not visible in some of the panels as they are smaller than the icons.

Supplementary Figure 3

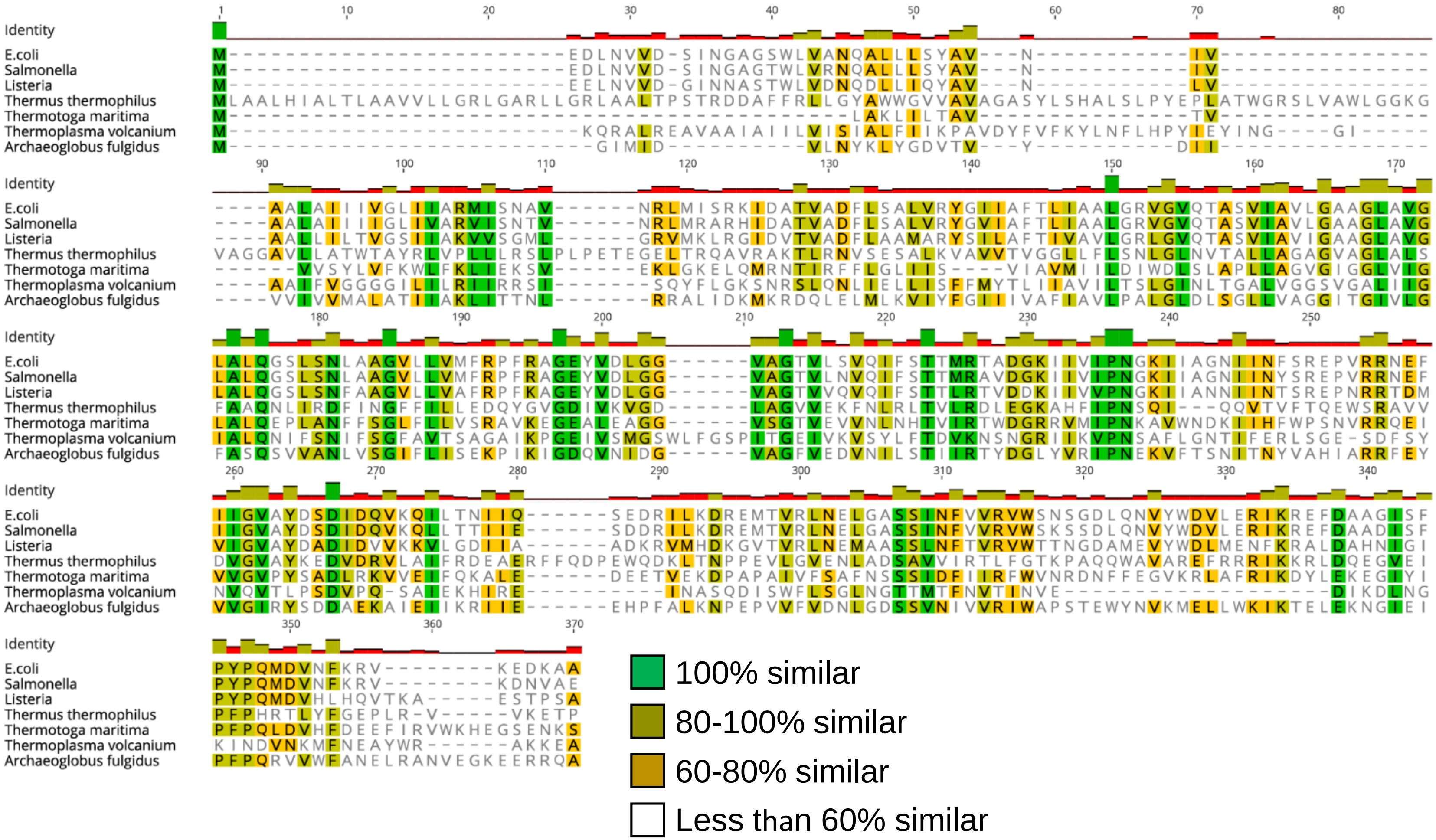


**Supplementary Figure S4. Sequence conservation of small mechanosensitive channels.** The sequence alignment of the MscS homologues tested in Figure 7 was performed using the Geneious Prime (2025.0.3) software, using the built-in MUSCLE alignment tool with default parameters.
